# A hormonally regulated gating mechanism controls EMT timing to ensure progenitor cell specification occurs prior to epithelial breakdown

**DOI:** 10.1101/2025.07.19.665116

**Authors:** Andrew T Plygawko, Jamie Adams, Zack Richards, Kyra Campbell

**Affiliations:** School of Biosciences, The University of Sheffield, S10 2TN, Sheffield, UK

## Abstract

Epithelial-to-mesenchymal transitions (EMTs) are essential for morphogenesis, converting static epithelial cells into migratory mesenchymal cells. As EMT disrupts epithelial barriers, its timing must be tightly regulated during development. We investigated this regulation in the *Drosophila* posterior embryonic midgut, where the GATA transcription factor Serpent (Srp) drives a collective partial EMT in the principal midgut epithelial cells (PMECs). Srp is expressed well before PMEC-EMT initiation and is also required for an earlier event: the specification and delamination of endoblasts, which give rise to different midgut lineages including adult midgut progenitors (AMPs). Here we identify the steroid hormone ecdysone as a temporal cue that triggers PMEC-EMT. Although Srp activates and represses numerous genes during early stages of midgut development, including polarity regulators such as *crumbs (crb),* the removal of polarity proteins and E-Cadherin from the membrane of PMECs only properly occurs in the presence of ecdysone signalling. This suggests that Srp primes cells transcriptionally for EMT, which proceeds only once polarity and adhesion proteins are downregulated post-translationally. When PMECs undergo a premature EMT, this leads to a reduction in the number of progenitor cells, suggesting that maintaining epithelial integrity until after endoblasts are selected is crucial for early fate decisions. These findings reveal that ecdysone functions as a EMT gatekeeper, ensuring that EMT is temporally restricted to occur only after progenitor specification, and in coordination with other developmental processes. More broadly, our work shows that in this context, transcriptional priming of EMT can be uncoupled from its execution, which is post-translationally triggered by hormonal signals. Given the conserved role of nuclear steroid hormone receptors as GATA co-factors and their association with poor prognosis in cancer, this may represent a broader strategy by which systemic cues regulate EMT in development and cancer.

## Introduction

Epithelial cells are essential building blocks of animal tissues, forming tightly connected sheets that provide both structural support and protective barriers roles during development and in adult organs. The regulated transformation of epithelial cells into mesenchymal cells, a process known as epithelial-to-mesenchymal transition (EMT), is essential for embryonic development^1^. EMT enables cells to delaminate from epithelial sheets, migrate, and build organs and tissues throughout the body. Given the critical barrier functions of epithelia, EMT must be tightly controlled both spatially and temporally. Premature or dysregulated EMT can lead to developmental defects and has been implicated in pathological conditions such as cancer metastasis and fibrosis^2^.

A key feature of developmental EMTs is the presence of EMT-inducing transcription factors (EMT-TFs)^3^, which are often expressed well before any overt cellular changes^4–6^. For example, the EMT-TF Snail is expressed very early in the presumptive *Drosophila* mesoderm^7^. Although Snail is transcriptionally active at this stage, with known targets already being expressed^8–10^, EMT does not occur until after the mesoderm has been internalised and the actomyosin tension released^4,11^. Studies in vertebrates have provided further evidence that the activity of EMT-TFs can be modulated by the mechanical properties of the cellular environment, highlighting a conserved role for mechanical cues in regulating EMT timing^4,5,12^. However, additional mechanisms are likely required to regulate the precise timing of EMT onset and ensure that EMT only proceeds once appropriate developmental milestones are reached. How EMT-TF activity is temporally regulated beyond mechanical influences remains poorly understood.

The embryonic midgut of *Drosophila melanogaster* presents a well-characterised and experimentally accessible system to study how the timing of EMT is developmentally regulated^13–15^. The midgut originates from two epithelial invaginations positioned at opposite poles of the blastoderm-stage embryo^16,17^. During gastrulation, specific cells within these anterior and posterior invaginations - the principal midgut epithelial cells (PMECs) - undergo a collective, partial EMT^13,18^. They temporarily adopt mesenchymal behaviour, migrate towards each other and subsequently undergo a mesenchymal-to-epithelial transition, and form the epithelial lining of the embryonic gut tube^14,19^.

We have recently shown that prior to EMT, another key developmental event occurs^15^. Individual cells, termed endoblasts, delaminate apically into the lumen of each midgut primordium. These transient cells undergo a single asymmetric cell division to produce either an enteroendocrine cell (EE) or an adult midgut progenitor cell (AMP). A small subset of endoblasts in the distal tip of the posterior midgut instead generates EEs or interstitial cell precursors (ICPs). These populations of endoblast-derived cells collectively migrate with the PMECs and contribute to the formation of cell lineages required for later developmental stages. In particular, the AMPs form progenitor cells which subsequently give rise to all cells in the adult midgut epithelium, including the intestinal stem cells^20,21^.

EMT in the posterior midgut is driven by the GATA transcription factor Serpent (Srp). Srp shows many parallels with Sna – both are required for EMT in their respective tissues, but also have earlier roles in determining cell fate, with Srp necessary for endoderm specification, and Sna for mesoderm^7,22,23^. Previous work has shown that Srp is first expressed during stages 5-6 of development, yet EMT does not occur until roughly 2 hours later^13^. This is not simply due to a delay in Serpent activity as *hindsight* (*hnt*), a known Srp target, is already expressed during stage 6 (Fig. 1A,B and^24^). Furthermore, Srp is also required for the specification of AMP, EE, and ICP cell types, processes that precede its role in EMT^25^. These observations raise a fundamental question - given the early expression and activity of Srp, what prevents premature EMT?

**Figure 1.**
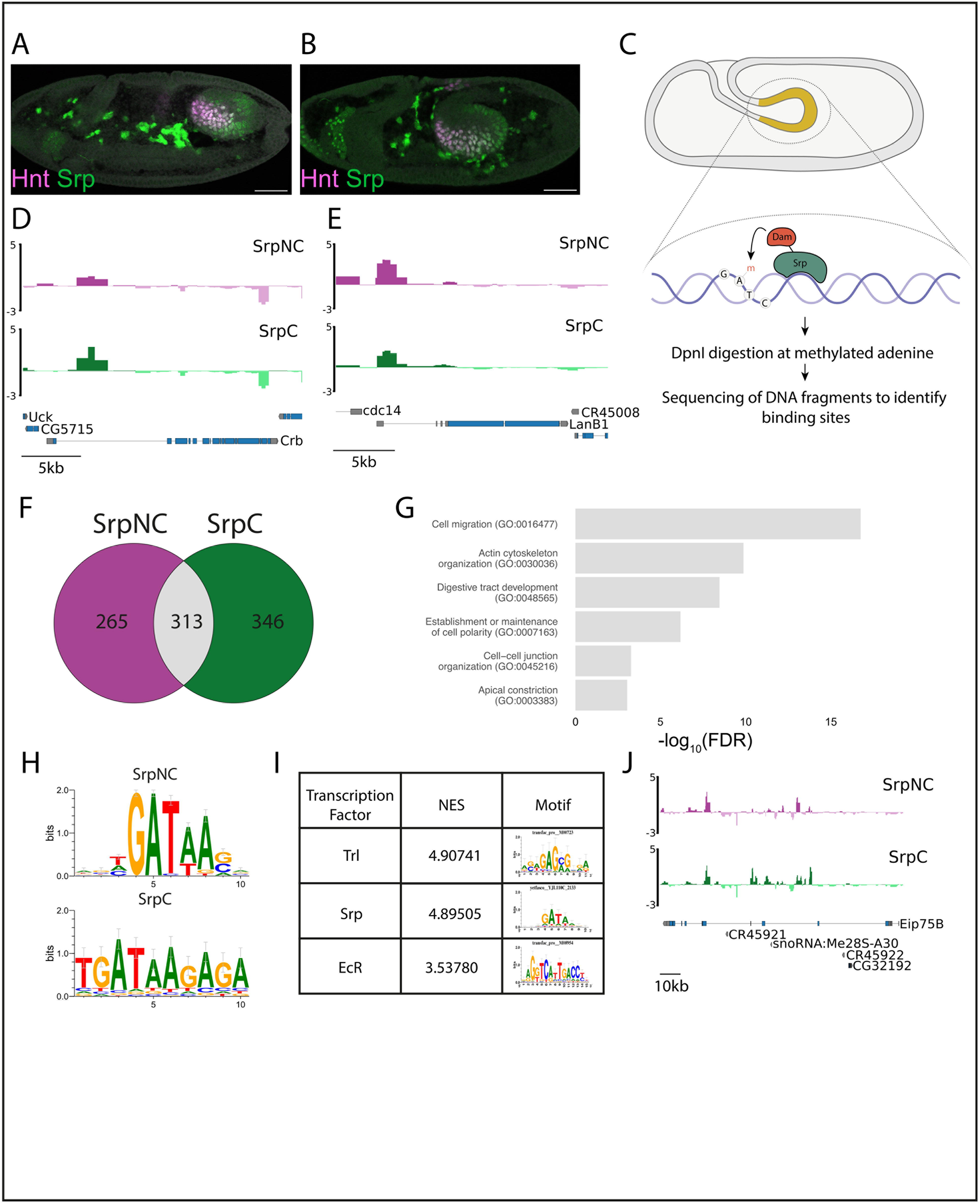
Serpent binding sites during Midgut-EMT overlap with the Ecdysone Receptor binding motifs. (A,B) Expression of Srp (green) and Hnt (magenta) in stage 6 (A) and stage 7 embryos. (C) Schematic of TaDa for Srp-Dam. (D,E) Srp binding sites associated with *crb* (D) and *LanB1* (E). The y-axis is a log_2_ score for Srp binding enrichment over background binding. (F) Venn diagram depicting the overlap between SrpNC and SrpC target genes. (G) Graphical representation of the overrepresented GO annotation classes from shared SrpNC and Srp targets. (H) GATA motifs are enriched in SrpNC and SrpC targets. (I) Motif enrichment analysis of shared SrpNC and SrpC targets revealed enrichment for the position weight natrices (PWMs) for Trithorax-like, Srp and EcR showing enriched binding motifs in shared SrpNC and SrpC targets. (J) Srp binding sites associated with the EcR target Eip75B. Scale bars, 50 μm (A,B).

Here, we uncover a role for the steroid hormone 20-hydroxyecdysone (ecdysone) in the temporal control of Srp-driven EMT. Given that GATA factor activity is often regulated through interactions with transcriptional co-factors^26^, we generated a genome-wide binding profile for Serpent in the endoderm and searched for common binding motifs to identify putative co- regulators. This analysis revealed that Serpent transcriptional targets can be subdivided by the presence of ecdysone-responsive motifs. Although Serpent alone regulates a broad set of target genes, including those involved in epithelial polarity such as *crumbs* (*crb*), timely cellular de-adhesion and EMT occur only in the presence of ecdysone signalling. Notably, only PMECs respond to ecdysone, whereas endoblasts, which delaminate earlier, do not. Furthermore, when PMECs undergo a premature EMT, this leads to a reduction in the number of progenitor cells, suggesting that maintaining epithelial integrity until after endoblasts are selected is crucial for early fate decisions. These findings reveal that ecdysone functions as a temporal gate for EMT, ensuring that breakdown of the midgut epithelium only occurs after progenitor specification is complete. Given the strong conservation of hormone-responsive transcriptional machinery, and the known cooperation between nuclear hormone receptors and GATA factors in mammalian systems^27–29^, we propose that hormone gating may act as a conserved mechanism for the temporal regulation of EMT onset.

## Results

### Mapping Serpent binding sites during midgut EMT

Srp is a GATA transcription factor, part of a highly conserved family that regulates diverse developmental processes in a context-dependent manner^26^. GATA factor activity is often modulated by interactions with other transcription factors, and their ability to activate or repress specific target genes depends on the presence of appropriate co-regulators. These interactions help confer spatial or temporal specificity, which is essential for carrying out distinct functions, as GATA factors often have multiple roles despite being broadly and constitutively expressed within developing tissues^30^. Given that Srp is expressed well before the onset of EMT in the midgut, we hypothesised that additional factors may be necessary to enable Srp to induce EMT. Specifically, we reasoned that Srp may function in combination with other transcription factors to regulate distinct aspects of endoderm specification and EMT onset. To identify such co-regulators, we aimed to generate a genome-wide binding profile for Srp in the endoderm, spanning the period before and after EMT initiation (Stages 8-11).

To achieve this, we employed Targeted DamID (TaDa), a cell type-specific profiling method based on DamID^31^ that enables the detection of protein-DNA interactions *in vivo* without the need for cell isolation, fixation, or specific antibodies^32,33^ (Fig. 1C). While GATA factors are characterised by the presence of two highly conserved zinc fingers, Srp is either produced in an isoform that contains both zinc fingers (SrpNC) or just one zinc finger (SrpC)^34^. While Srp isoforms differ in the capacity to activate certain target genes *in vivo,* analyses of isoform specific loss-of-function mutants suggests that either is sufficient to drive EMT during midgut formation^35^. We therefore generated two constructs, UAS-Dam-SrpNC and UAS-Dam-SrpC (see Methods), and reasoned that *bona fide* EMT targets would be bound by both isoforms. This strategy allowed us to control for isoform-specific or non-specific background binding.

Analysis of peaks of binding indicate that both SrpC and SrpNC primarily bind to intergenic regions and in sites overlapping with the transcription start site (Supp. Fig. 1). This shows parallels with vertebrate orthologs of Srp, GATAs 4 and 6, which are similarly expressed in the endoderm and are also capable of driving an EMT^13,36,37^. GATA4 sites are mainly intragenic (primarily in the first intron) and intergenic regions, with only a small subset of the binding sites targeting upstream promoters, while GATA6 binds to promoters, intragenic regions and distal intergenic regions (reviewed in^30^).

We identified 578 target genes bound by SrpNC and 659 for SrpC, with 313 of these targets shared between the two isoforms (Fig. 1F). Supporting the validity of our analysis, we replicated binding peaks previously identified by Chromatin Immunoprecipitation followed by Polymerase Chain Reaction (ChIP-PCR) in the first intron of *crumbs* (*crb), Laminin B1*, and *Laminin B2*, as well as identifying binding in the known Srp target GATAe (Fig. 1D,E, Supp. Fig 2 and^13,38^). Furthermore, the most common motif found among genes bound by either isoform was the consensus WGATAR sequence^39^ (Fig 1H). To further characterise common binding sites, we used established pipelines^40^ to extract the associated genes and carried out gene set enrichment analysis. Overrepresented Gene Ontology (GO) terms included categories related to cell migration, actin cytoskeleton organisation, digestive tract development, establishment of maintenance of cell polarity, cell-cell junction organisation and apical constriction (Fig. 1G), reflecting the multifaceted roles of Srp in midgut morphogenesis and EMT.

### Ecdysone signalling is required for EMT in the midgut

To identify putative co-factors associating with Srp when binding to the genome, we searched for transcription factor-binding motifs that are overrepresented in the genes bound by both Srp isoforms using i-*cis*Target^41^. Top hits included Serpent itself, the GAGA factor Trithorax-like (Trl) and notably, the Ecdysone Receptor (EcR) (Fig. 1I). In line with EcR being a potential co- factor, we found Srp binding sites in the first intron of the known ecdysone-responsive gene *Eip75B* (Fig. 1J). These findings raised the possibility that ecdysone signalling may cooperate with Srp to regulate target gene expression during early midgut development.

Given the well-established role of hormones in regulating developmental timing across species (reviewed in^42^), the enrichment of EcR motifs within Srp-bound chromatin regions was particularly striking. EcR is the nuclear receptor for the steroid hormone ecdysone, which coordinates major developmental transitions in *Drosophila*, including molting and metamorphosis^43^. While a requirement for ecdysone in midgut morphogenesis during mid- embryogenesis has been reported^44,45^, its presence in Srp-bound chromatin regions during stages 8–11 of embryogenesis suggests a previously unappreciated role for ecdysone signalling in regulating earlier stages of midgut development.

To test whether Edysone signalling contributes to midgut-EMT, we blocked the functioning of endogenous EcR by expressing a dominant negative form of the EcR (UAS-EcR-DN) together with the membrane marker UAS-SrcGFP, using hkb-Gal4, which expresses in midgut cells from very early stages of development^13^. At the stage when midgut cells have normally rounded up and multilayered (Fig. 2A, arrows), we observed that midgut cells with disrupted ecdysone signalling retained a more columnar, epithelial-like morphology (Fig. 2B, arrows).

**Figure 2.**
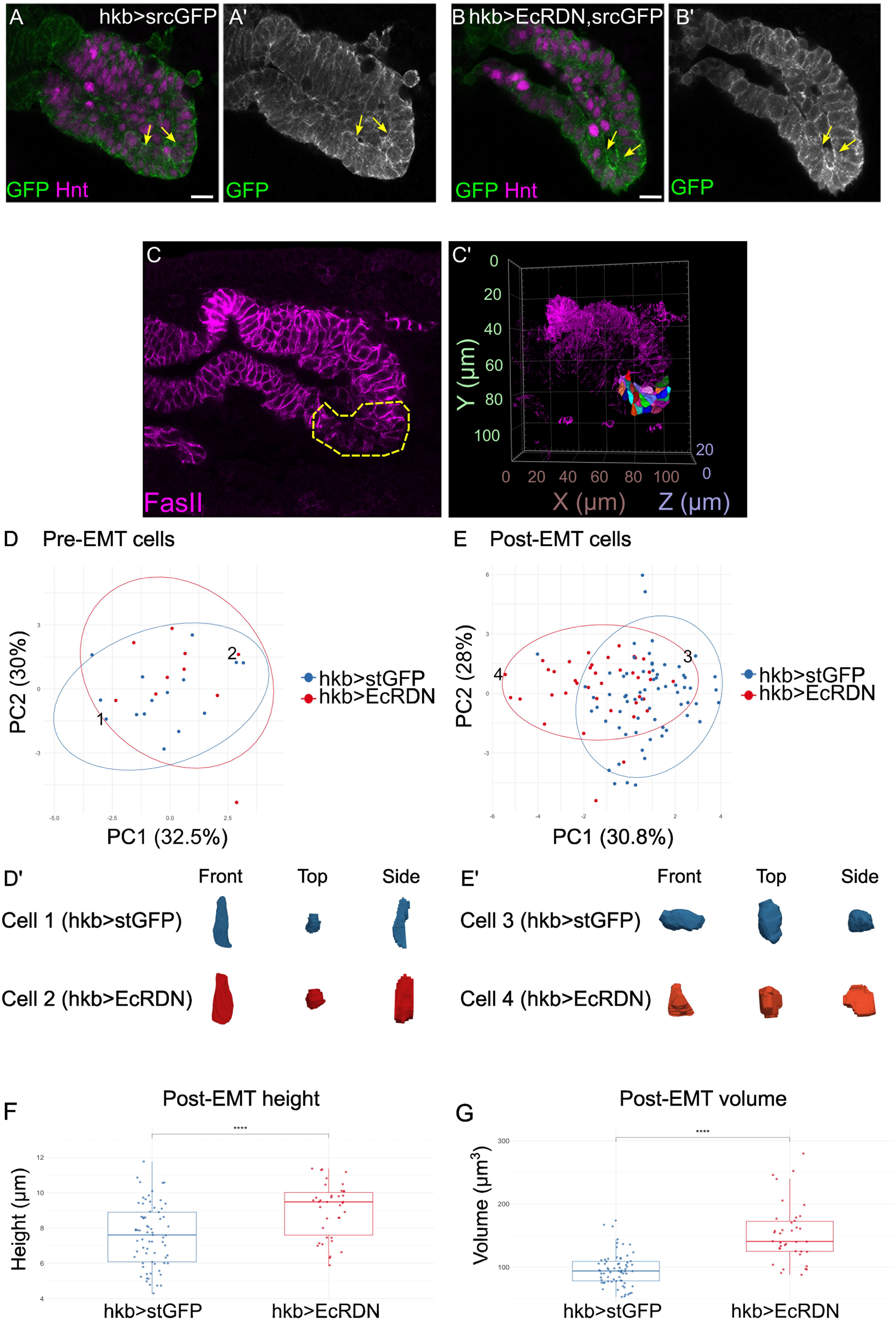
Ecdysone Receptor activity is required for changes in cell morphology that occur during EMT. (A,B) Immunofluorescence for GFP (green) and Hnt (magenta) in control embryos expressing Src-GFP alone (A), or in embryos co-expressing UAS-EcrDN (B). Yellow arrows point to the apical surface of PMECs in the distal region of the midgut, which are the first to undergo EMT in wildtype. (C) Representative image of a developing midgut. Dotted yellow outlines the region in which cells were segmented. (D,E) Principal Component Analysis plot for cells segmented from hkb-stGFP (blue) or hkb-EcRDN (red) midguts at a Pre-EMT (D) or Post-EMT timepoint (E). Each dot represents a single cell. Ellipses represent 95% confidence regions for each genotype. Numbers indicate which cells have been selected to be highlighted in D’ and E’. (F, G) Boxplots for cell height and volume in each genotype. The box height represents interquartile range, with asterisks showing significant differences via t- test, **** is *p*<0.0001. Scale bars, 10 μm (A,B).

To characterise these shape differences in greater detail, we used single cell morphometric analysis (scMorphometrics), a powerful approach for quantifying cell shape in 3D from confocal image stacks^46^. scMorphometrics allows an unbiased measurement of cell morphology, which can then be projected onto 2D shape trajectories^46,47^.

We segmented cells from 3D confocal z-stacks in wild type embryos at pre and post-EMT timepoints, focusing on the more distal portion of the midgut (Fig. 2C, C’), and then segmented cells from the same region of the midgut, and at the same stages, from embryos expressing EcR-DN. A principal component analysis of cellular features revealed that pre-EMT, both wildtype and midgut cells expressing EcR-DN are indistinguishable in their cell shape, with all cells exhibiting the elongated, columnar shape characteristic of epithelial cells (Fig. 2D). In contrast, post-EMT, cells defective for ecdysone signalling appear to possess a distinct morphology from their wild type counterparts (Fig. 2E). Principal Component 1 is comprised of metrics related to cell size (Supp. Fig. 3A), while Principal Component 2 describes the regularity of the cell shape (Supp. Fig. 3B). The negative contribution of maximal lateral width to PC2 (Supp. Fig. 3C) suggests that cells widen as their shape becomes more irregular, perhaps reflecting a loosening of tension at the onset of EMT. EcR-DN expressing cells score higher in both Principal Components, meaning these cells are taller and larger than in wild type (Fig 2E, F, G). Furthermore, their high score in PC2 suggests that EcR-DN cells possess a more regular surface which may be indicative of tighter packing within the tissue. Indeed, these cells are more columnar, often with clear apical surfaces, and fail to adopt the wider and shorter mesenchymal morphology seen in wild type midguts (Fig. 2E). The fact that apical domains remain more established in EcR-DN cells suggests that a key step in midgut-EMT is the constriction and removal of the apical surface.

### Ecdysone signalling is required for post-translational downregulation of apicobasal/junctional proteins from the plasma membrane

We previously showed that a key component of Srp-driven EMT is the direct transcriptional repression of the apicobasal polarity determinant *crumb (crb)*, which leads to a loss of epithelial polarity^13^. In contrast, junctional proteins such as dE-Cadherin (dE-Cad) and Bazooka ((Baz), *Drosophila* Par-3) are not transcriptionally repressed. Instead, these proteins are removed from the cell membrane during EMT^13^ and are later recycled to mediate dynamic adhesions as cells undergo collective migration^18^.

To determine which aspects of EMT require ecdysone signalling, we first examined *crb* RNA expression dynamics in the developing midgut using fluorescence *in situ* hybridisation (FISH)^48^. We found that *crb* expression is downregulated by stage 7, well before the onset of EMT (Fig. 3A-C). In EcR-DN embryos, *crb* repression occurs normally, indicating that Srp- mediated transcriptional repression of *crb* is independent of ecdysone signalling (Fig. 3D-F).

**Figure 3.**
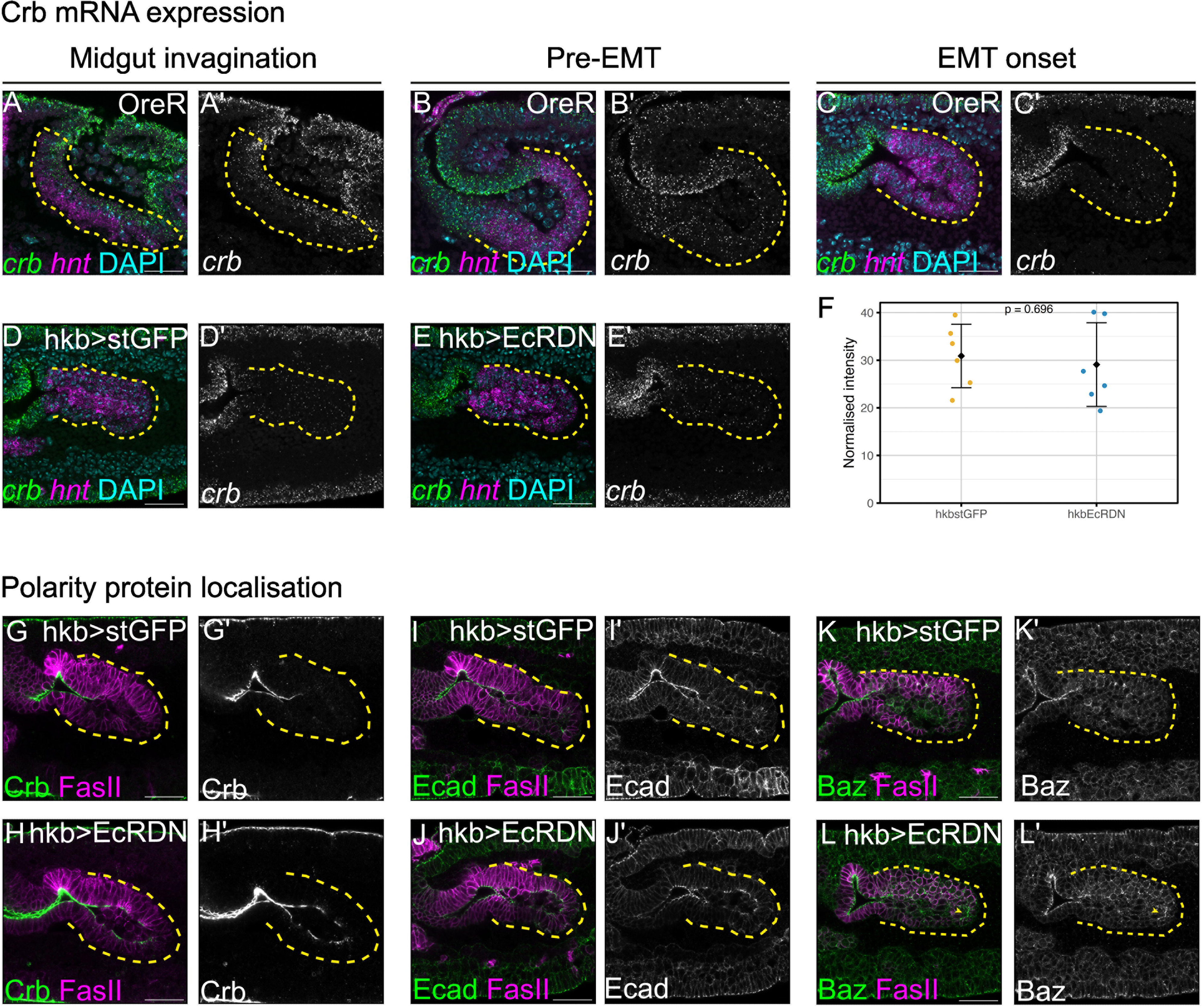
Ecdysone signalling is required for the downregulation of polarity and adhesion proteins from the cell membrane. (A-C) FISH for *crb* mRNA (green) and mRNA for the midgut marker *hnt* (magenta) in wild type embryos at different stages of development. (D,E) FISH for *crb* (green) and *hnt* (magenta) in hkb-stGFP (D), or hkb-EcRDN (E) stage 10 midguts. (F) Normalised fluorescence intensity measurements of *crb* RNA in stage 10 midguts in in hkb-stGFP and hkb-EcRDN conditions reveal no significant differences in RNA levels. Average values for each embryo were used to calculate the overall mean ± standard deviation, with p=0.696 calculated via t-test. (G, H) Immunofluorescent staining of stage 10 embryos stained for Crb (green) and FasII (magenta) in hkb-stGFP (G, G’) and hkb-EcRDN (H, H’) midguts. (I, J) Immunofluorescent staining of stage 10 embryos stained for Ecad (green) and FasII (magenta) in hkb-stGFP (I) and hkb-EcRDN (J) midguts. (K, L) Immunofluorescent staining of stage 10 embryos stained for Baz (green) and FasII (magenta) in hkb-stGFP (K) and hkb-EcRDN (L) midguts. Yellow dotted lines outline the posterior midgut in each image. Scale bars, 25 μm.

We next examined Crb protein localisation. Despite the early loss of *crb* RNA, Crb protein persists at the apical membrane of midgut cells until stage 10, when it is rapidly lost (Fig. 3G). In embryos with impaired ecdysone signalling, Crb is not downregulated from the membrane and remains apically localised (Fig. 3H). Similarly, dE-Cad and Baz also fail to be removed from the membrane when EcRDN is expressed in midgut cells (Fig. 3I-L).

Together, these data suggest that midgut cells are transcriptionally primed for EMT through early repression of *crb*, but loss of polarity and adhesion requires the ecdysone-dependent removal of Crb and junctional proteins from the membrane. Notably, in *crb* mutant embryos, EMT occurs prematurely in the midgut^13^, further supporting the idea that membrane-localized Crb maintains cells in an epithelial state until these proteins are actively cleared.

### Crb protein removal depends on different mechanisms in PMECs vs endoblasts

This led us to investigate how Crb is removed from the cell membrane. Previous studies have shown that binding of the E3 ubiquitin ligase Neuralized (Neur) to the Crb complex protein Stardust (Sdt) promotes Crb endocytosis^49,50^. We therefore hypothesised that ecdysone signalling may act upstream of Neur, triggering its expression to initiate membrane removal of Crb and the subsequent downregulation of junctional proteins at the appropriate developmental stage.

To test this, we first examined the spatial and temporal dynamics of *neur* expression in the posterior midgut. FISH revealed that *neur* is expressed in stage 9 embryos (Fig. 4A). However, rather than being expressed throughout the epithelium, which would be expected if it were involved in midgut EMT, it appears to be expressed in a salt and pepper pattern. As this is reminiscent of the localisation of endoblasts prior to delamination (Fig. 4C), we performed FISH for the endoblast marker *sna* and found that it co-localises with *neur,* both in early stages where endoblasts are still part of the epithelium, and later after the cells have moved into the midgut lumen (Fig. 4A,B).

**Figure 4.**
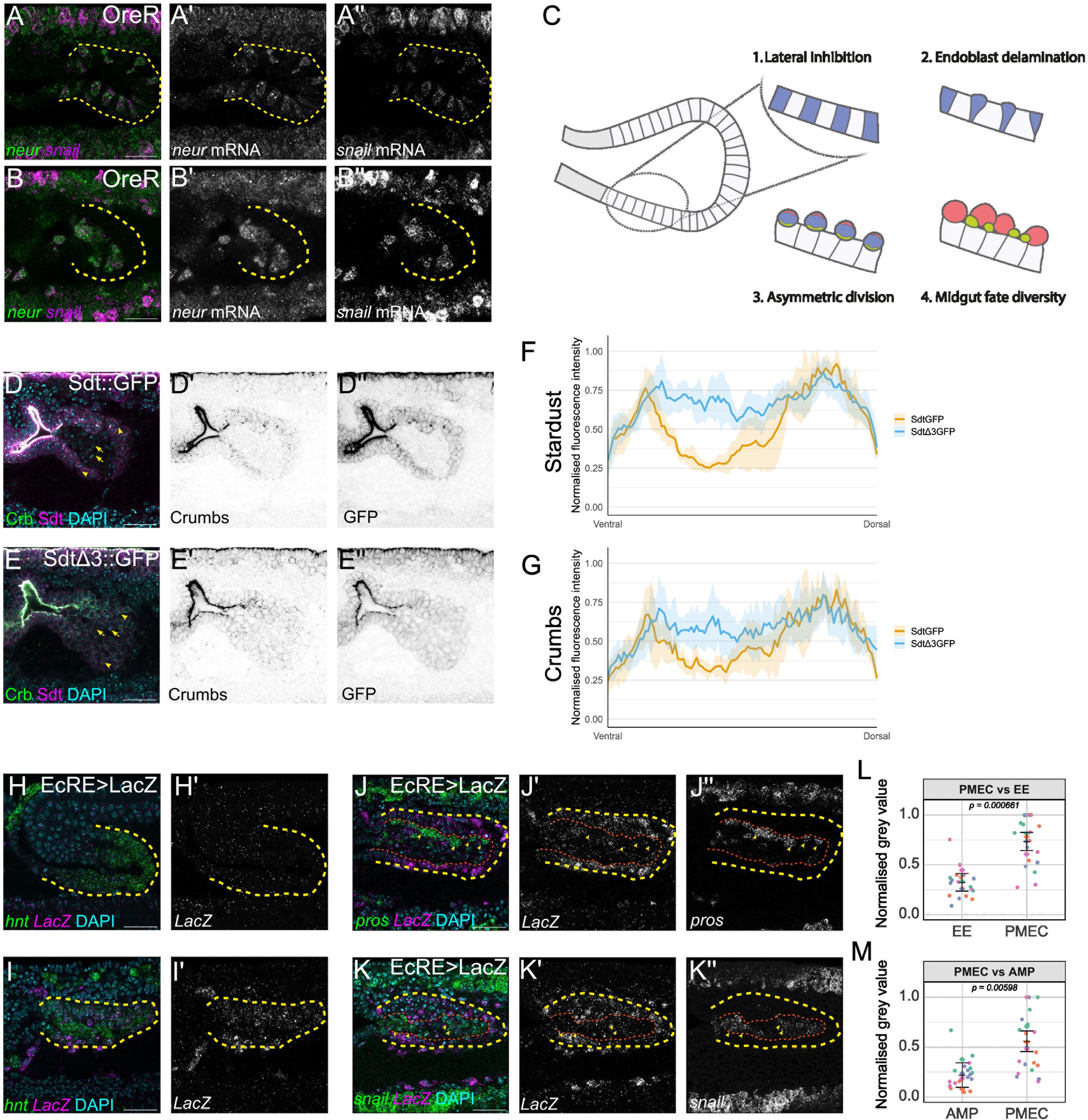
Different methods of Crb protein removal are used in different midgut cell populations. (A, B) FISH for *neur* mRNA (green) and *snail* mRNA (magenta) in stage 9 (A) and stage 10 (B) midguts. (C) Cartoon of the steps leading to endoblast delamination. 1. Lateral inhibition results in differential Notch-Delta activity in neighbouring cells. 2. Endoblasts are selected by low Notch activity and delaminate apically. 3. Endoblasts asymmetrically divide. 4. Differential inheritance of cell fate determinants produces different cell fates. (D, E) Immunostaining for Crb (green) and GFP-tagged Stardust (magenta) with (D, Sdt::GFP) or without (E, Sdt.3::GFP) the Sdt-Neur interaction site. Yellow arrows point to cells within the endoblast-derived population, while arrowheads point to cells in the PMEC layer. (F,G) Normalised fluorescence intensity profiles of GFP-tagged Stardust (F) and Crumbs (G) across the midgut, showing a decrease in intensity from endoblast-derived cells in the wildtype Sdt::GFP embryos, which and a more uniform expression in Sdt.3::GFP embryos, which lack the Sdt-Neur interaction. (H, I) FISH for midgut marker *hnt* mRNA (green) and *lacz* mRNA (magenta) where *lacz* is driven under control of the ecdysone response element in stage 7 (H, H’) and stage 10 (I) embryos. (J, K) FISH for *lacz* mRNA and the EE marker *pros* (J) or the AMP marker *snail* (K) in stage 10 embryos. Yellow dotted lines show the midgut, while orange dotted lines show the boundary between PMECs and AMPs/EEs. Arrows point to individual EEs (J) and AMPs (K). Fluorescence intensity of *lacz* signal was measured for different cell types across multiple embryos, as laid out in super plot format (L, M). Average values for each embryo were used to calculate the overall mean ± standard deviation, with p-values calculated via t-test. Scale bars, 25 μm.

This led us to ask whether different mechanisms remove Crb in the different midgut cell populations. Because Neur is also a key component of Notch signalling, which is required for endoblast selection, direct genetic perturbation of *neur* would likely interfere with endoblast formation and specification of the AMPs and EEs. Instead, we used a more targeted approach by analysing embryos carrying the *sdtΔ3:GFP* allele, in which the Neur-interaction domain of Std is specifically disrupted^50^. This mutation abrogates Neur-mediated endocytosis of Crb without affecting other Sdt-dependent functions.

In control embryos, Crb and Sdt were completely cleared from endoblast cells and their progeny (Fig. 4D, arrows), while low levels remained on the membrane and intracellularly in the PMECS (Fig. 4D, arrowheads). In contrast, in *sdtΔ3:GFP* mutants endoblasts, EEs and AMPs retained both Crb and Sdt (Fig. 4D-G), indicating that the interaction between Sdt and Neur is essential for Crb removal in these cells. However, the absence of *neur* expression in PMECs suggests that a different, as-yet unidentified, pathway must traffic Crb from the membrane in these cells. Importantly, given the persistence of Crb in PMECs when ecdysone signalling is blocked, this suggests that the alternate pathway operates under hormonal control.

### Ecdysone signalling is active only in PMECs and is dispensable for progenitor cell specification

The expression of Neur suggested a previously unrecognised cellular specificity to the remodelling events occurring around the time of endoblast delamination and midgut EMT. To determine if there is any specificity to ecdysone signalling, we next examined the spatial distribution of ecdysone pathway activity using an ecdysone response element (EcRE)- driven *LacZ* (*EcRE-LacZ)* reporter^51^. FISH for *lacZ* showed that there is no ecdysone activity during early stages of midgut development (Fig. 4H), and it is first detected in the PMECs during stages 10–11, corresponding with the onset of EMT (Fig 4I-K, region between dotted yellow and red lines denote the PMECs, quantified in L,M). In contrast, endoblasts and their progeny showed very little detectable *lacZ* expression at any stage examined (Fig. 4j-K, quantified in L,M). This suggests that ecdysone signalling is spatially restricted to the PMECs and is not active in the delaminating endoblasts.

To confirm that there is no role for ecdysone in endoblasts, we next asked whether blocking ecdysone signalling affected the formation of their daughter cells – the EEs and AMPs. Cell counts revealed no significant differences in the number or AMPs and EEs compared to wild-type controls (Supp. Fig. 4), indicating that ecdysone signalling is dispensable for endoblast delamination and EE and AMP cell specification.

Together, this points to two temporally separable morphogenetic events in the early midgut – endoblast selection and delamination from the epithelial midgut primordium, followed later by EMT in the outer epithelial PMECs. The timing of EMT onset in PMECs appears to be triggered by the downregulation of Crb protein, and the subsequent removal of junctional proteins from the cell membrane, which is under ecdysone control.

### Ecdysone controls the timing of PMEC-EMT in a Srp-dependent manner

Our findings thus far strongly indicate that ecdysone signalling acts as a key trigger to initiate PMEC-EMT by acting as a co-regulator for a subset of Srp targets. Srp is present in the embryo from as early as stage 6. Therefore, if the temporal control of EMT is governed by the timing of exposure to the ecdysone hormone, this would suggest that exposing posterior midgut cells to ecdysone at earlier stages should trigger a premature EMT.

To test this, we cultured embryos with exogenous 20-hyroxyecdysone (20E), the physiologically active form of the hormone. Previous studies have demonstrated that 20E treatment can induce premature and widespread activation of ecdysone signalling across the embryo^45^. In untreated control embryos at stage 7, the posterior midgut epithelium remains fully intact and epithelial, as indicated by tight apical localisation of Baz, which we used as a readout for polarity (Fig. 5A-A1’). When stage 7 embryos are exposed to 20E, the midgut appears convoluted, with ectopic folds, characteristic of apically constricted epithelial cells, and there are some signs of disrupted polarity, suggesting that midgut cells are prematurely initiating aspects of EMT (Fig. 5B-B1’). Together with the loss-of-function experiments, these findings support a model in which ecdysone controls the timing of EMT in the posterior midgut.

**Figure 5.**
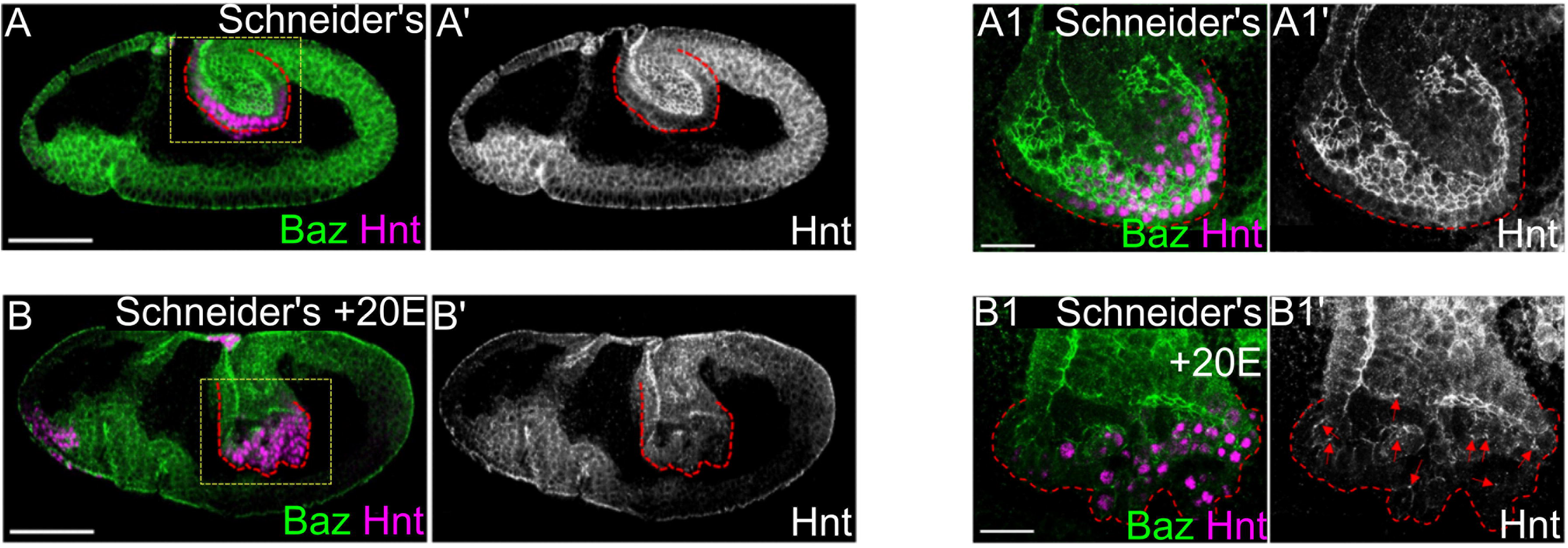
Early ecdysone exposure triggers a premature midgut-EMT in a Srp- dependent manner. Immunostaining of wild type embryos for Baz (green) and Hnt (magenta) in stage 7 embryos bathed in Schneider’s Medium (A-A1’) or Schneider’s Medium supplemented with 20E (B-B1’). The red dotted lines outline the midgut of each embryo, while the yellow box indicates the region highlighted in A1 and B1. Scale bars = 50μm.

### Premature PMEC-EMT leads to a reduction in progenitor cell specification

Having established that ecdysone signalling regulates the timing of PMEC-EMT onset, we next sought to understand why the temporal control of EMT is so critical for proper midgut development. Notch signalling is known to play an important role in early fate decisions in the midgut, acting upstream of the specification of endoblasts, EEs and AMPs. Given the known importance for both topological tissue organisation and for Crb interactions in Notch signalling^52,53^, we wondered if maintenance of an epithelium is required until Notch signalling has selected endoblasts. We reasoned that premature EMT, by disrupting epithelial integrity and reducing cell-cell contacts, might negatively affect endoblast selection and progenitor specification.

To investigate this, we took advantage of our previous finding that in *crb* mutants there is a premature EMT^13^. We therefore immunostained *crb* mutants for markers for EEs and AMPs and performed cell counts. We found that in this condition, the numbers of AMPs and EEs are greatly reduced (Fig. 6). This suggests that maintaining the epithelial sheet during the window of endoblast selection and progenitor specification is critical to ensure appropriate levels of Notch signalling takes place.

**Figure 6.**
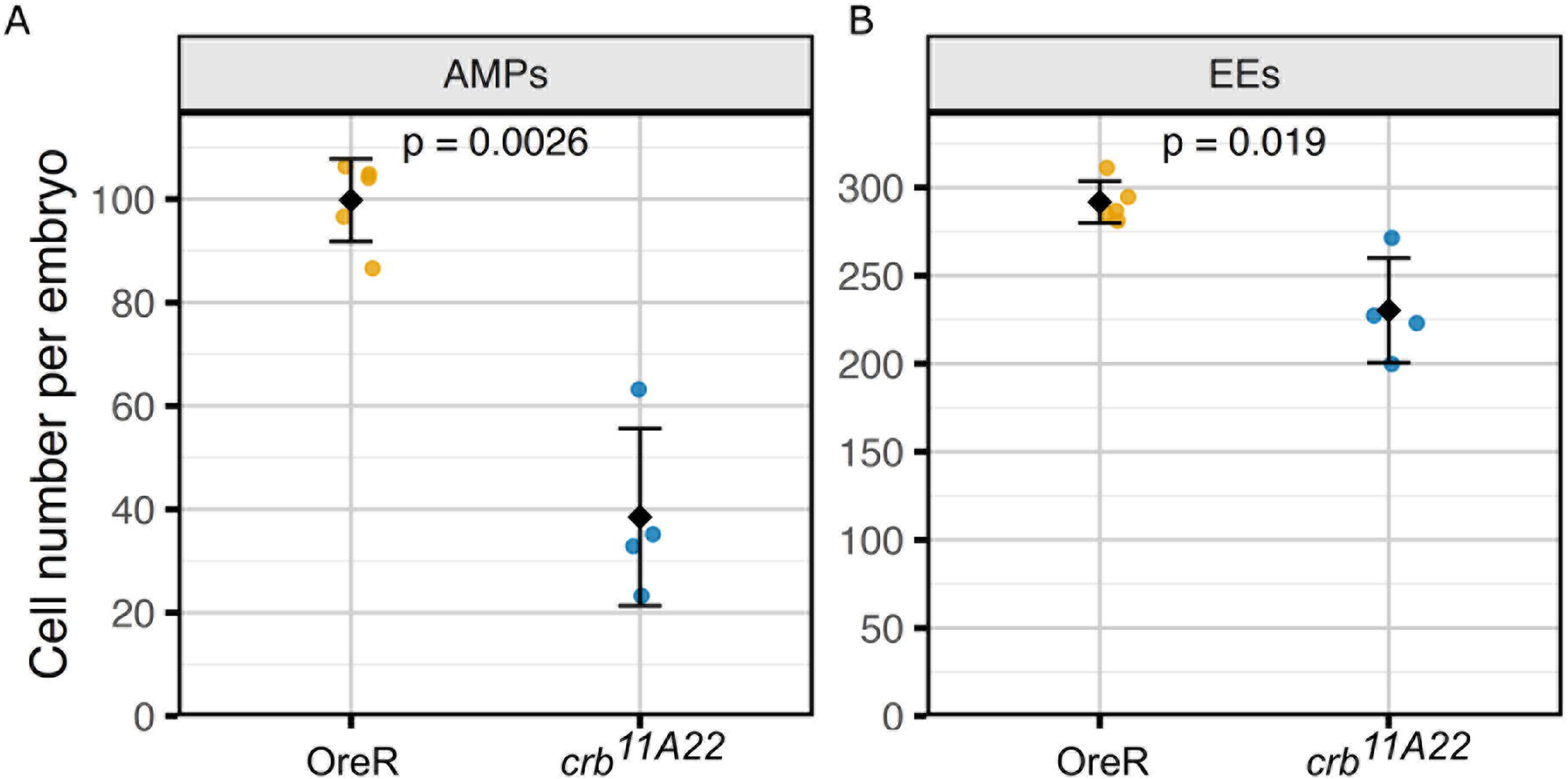
Premature loss of cell polarity reduces the number of EEs and AMPs. Cell counts in stage 15 wild type and *crb* mutant embryos for AMPs (A) and EEs (B). Each dot represents the cell count from an individual embryo. Black diamonds represent the mean with error bars for standard deviation, with p-values calculated via t-test.

## Discussion

In this study, we show that the steroid hormone ecdysone plays an essential role in the temporal control of EMT during *Drosophila* midgut development. While the GATA factor Srp initiates transcriptional priming by directly repressing *crb* early in development, we find that physical changes associated with EMT—such as junctional protein removal and loss of polarity—are delayed and require ecdysone signalling. This temporal separation between transcriptional priming and execution ensures that EMT does not proceed prematurely, allowing other developmental processes, such as cell fate specification via Notch mediated lateral inhibition, to occur in a properly organised epithelium.

Intriguingly, our findings suggest that two temporally and mechanistically distinct types of epithelial remodelling events occur in close succession in the midgut: endoblast delamination and PMEC-EMT. While endoblast delamination has not previously been classified as an EMT, it shares key morphological features with it—including junctional detachment, loss of apicobasal polarity and exit of cells from the epithelial layer into the midgut lumen. Our data suggest that, despite occurring within the same tissue, endoblasts and PMECs undergo two distinct partial EMTs. Endoblasts express Sna, individual delamination initiates with a basal constriction and cells downregulate Crb via the ubiquitin ligase Neur. In contrast, PMEC-EMT is triggered later, occurs collectively and involves apical domain remodelling, likely via constriction. It requires both Srp activity and ecdysone signalling and proceeds via the Neur- independent removal of polarity and adhesion proteins from the plasma membrane. This highlights that observed mechanistic and morphological differences in EMT are not merely context-dependent, but that even within a single epithelial tissue, at least two distinct EMT mechanisms can be deployed.

A key mechanistic insight from this study is that EMT in the midgut is triggered not simply by transcriptional changes, but by the removal of polarity and junctional proteins from the plasma membrane. While the transcriptional repression of adhesion and polarity factors like E-Cad and Crb have long been considered a hallmark of EMT, our data demonstrate that midgut cells retain apically-localised E-Cad and Crb protein for several hours after *crb* transcription is repressed. Only upon ecdysone signalling are Crb and other junctional proteins removed from the membrane, allowing cell shape changes and detachment. This two-step mechanism— transcriptional priming followed by hormone-triggered protein clearance—ensures that EMT occurs with high temporal precision.

Post-translational control of junctional disassembly has been observed in other EMT contexts, including in the *Drosophila* mesoderm, where adherens junction disassembly and internalisation occurs rapidly upon the release of actomyosin tension^4^. Our findings suggest that ecdysone signalling provides an additional route for post-translational EMT regulation, allowing transcriptionally primed cells to respond to systemic cues and remodel rapidly when the developmental context permits. It is possible that this mechanism represents a conserved feature of hormone-regulated EMTs. Indeed, ecdysone has recently been implicated in other *Drosophila* EMT and EMT-like contexts, including in the mesoderm^54^, follicle cell reorganisation^55^, and fat body remodelling^56^, although its molecular role in those processes remains unclear.

This work also provides insight into how EMT timing intersects with fate decisions. We have recently shown that progenitor cells arise from endoblasts, which are selected from the midgut epithelium via Notch–Delta–mediated lateral inhibition^15,57^. In this process, Notch signalling becomes inactive in the endoblasts, allowing them to delaminate into the midgut lumen, while it remains active in the neighbouring epithelial cells. Interestingly, we observe a reduction in progenitor cells in *crb* mutants, suggesting that the absence of Crb leads to overactivation of the Notch pathway in midgut cells. This could result directly from the loss of Crb protein at the apical membrane. In the developing pupal wing, for example, Crb is required for proper apical localization of Notch, and its absence leads to ligand-independent Notch activation and subsequent defects in cell fate specification^58^. Similarly, in zebrafish, Crb has been shown to interact with Notch in cell culture, and its overexpression reduces Notch activity^59^. Additionally, premature EMT in *crb* mutants may lead to early epithelial multilayering, which will alter the spatial context of Notch signalling. In a monolayered epithelium, cells interact laterally; in a multilayer, cells contact neighbours apically and basally as well, increasing the number of signalling interfaces and raising the possibility that multilayering leads to broader or ectopic activation of Notch. Thus, either through altered membrane organisation or tissue architecture, premature EMT disrupts Notch patterning and reduces progenitor output.

These findings highlight a previously unrecognized link between epithelial architecture, hormone signalling, and the developmental timing of EMT. While the roles of Crb and tissue organization in modulating Notch signalling are well established, our data show that these factors also constrain when EMT can occur without compromising fate decisions. Specifically, we demonstrate that EMT must be precisely delayed until after Notch-mediated progenitor selection has taken place. This highlights the broader developmental principle that tissue remodelling events such as EMT must be tightly coordinated with fate specification, and that this coordination can be achieved by separating transcriptional priming from post-translational execution.

More broadly, this work suggests that steroid hormones may play a general role in temporally gating EMTs during development. Transcription factors like Snail, Twist, Srp, and Zeb proteins are expressed across many contexts and often well before morphological transitions occur. Their pleiotropic activity suggests that additional layers of regulation—such as mechanical input or hormonal signalling—may be needed to fine-tune EMT onset. In vertebrates, the GATA factors GATA2 and GATA3 are known to interact with nuclear hormone receptors such as Androgen and Estrogen receptors^27,60^ and are implicated in EMT-related processes and cancer progression^26,61^. It is tempting to speculate that similar transcription factor–hormone partnerships could coordinate EMT timing across species.

## Material and Methods

### Fly stocks and maintenance

*Drosophila melanogaster* stocks and crosses were maintained and performed on cornmeal molasses food at 25°C. Details for all genotypes and transgenes can be found in flybase (http://flybase.org) or in references listed here. Unless otherwise noted, stocks were obtained from the Bloomington Stock centre. Stocks obtained from Bloomington *Drosophila* Stock Center (NIH P40OD018537) were used in this study. The following lines were used in this paper: Hkb-Gal4 (gifted by Helen Skaer), UAS-EcR-B1-DN (BDSC:6869), UAS-stingerGFP (BDSC 84277), 48Y-Gal4 (BDSC 4935), OreR (BDSC 5), *crb^11A^*^22^ (gift by Elisabeth Knust^62^), Stardust::GFP (BDSC 99581)^47^, Stardust^Δ^^3^::GFP (BDSC 99580)^47^.

Embryos driving UAS-stingerGFP or UAS-srcGFP in the midgut using either hkb-Gal4 or 48Y- Gal4 (BDSC 4935), or OreR were used as wildtype controls.

### *Drosophila* embryo fixation and immunofluorescence

Embryos for standard stainings were dechorionated using 50% bleach for 3 minutes, fixed in 4% PFA for 20 minutes, and then devitellinised with manual shaking. Embryos were permeabilised and blocked in PBS + 0.2% Triton X-100 (PBT) + 0.1% BSA for 2 hours. Primary antibodies were incubated overnight at 4°C, while secondary antibodies were incubated for 2 hours at room temperature. Primary antibodies used were: goat anti-GFP 1:500 (AB6673), rabbit anti-GFP 1:1000 (PABG1), rat anti-E-Cad 1:100 (DCAD2, DSHB), mouse anti-FasII 1:20 (1D4, DSHB) mouse anti-prospero 1:100 (MR1A, DSHB), guinea pig anti-Snail 1:1000^4^ (a gift from Mo Weng), mouse anti-Hindsight 1:20 (1G9), rabbit anti-Bazooka N-term 1:200^63^ (a gift from Andreas Wodarz), and rat anti-Serpent^13^ (1:500). Stainings for E-cad were performed on fresh embryos fixed for 10 minutes. DAPI was used as a DNA stain at a concentration of 1:250. Secondary antibodies, used at a concentration ranging from 1:100-200, were as follows: Donkey anti-Goat Alexa Fluor Plus 488 (A32814), Donkey anti-Rabbit Alexa Fluor Plus 488 (A32790), Donkey anti-Mouse Alexa Fluor Plus 555 (A32773), Donkey anti-Rat Alexa Fluor Plus 555 (A48270), Donkey anti-Rabbit Alexa Fluor Plus 555 (A32794), Donkey anti-Mouse Alexa Fluor Plus 647 (A32787), Goat anti-Guinea Pig Alexa Fluor 647 (A-21450) Unless otherwise stated, secondary antibodies were sourced from Thermo Fisher Scientific. Embryos were mounted in ProLong Glass Antifade Mountant (Thermo Scientific) or Fluoromount G (Invitrogen).

### HCR *in situ* hybridisation

HCR *in situ* hybridisation was performed using an adapted version of previously published protocols^48,64^. HCR v3.0 probes (*pros*-B1, *sna*-B4, *hnt*-B1, *hnt*-B2, *lacz*-B3, *neur*-B3, *crb*-B1) and hairpins (B1 546, B1 647, B2 546, B2 647, B3 647 and B4 647) were synthesised by Molecular Instruments. Following standard fixation, embryos were incubated in Probe Hybridization Buffer (Molecular Instruments) at 37°C for 30 minutes then hybridised with Probe Hybridisation Buffer (Molecular Instruments) containing HCR probes overnight at 37°C. Embryos were washed four times for 15 minutes each, using pre-warmed Probe Wash Buffer (Molecular Instruments) at 37°C. Embryos were then washed twice at room temperature in 5x SSC buffer, pre-amplified in Probe Amplification Buffer (Molecular Instruments) for 10 minutes, and then incubated overnight in Probe Amplification Buffer containing snap-cooled HCR hairpins at room temperature in the dark. Excess hairpins were removed with two 5 min washes with 5x SSC, followed by 2 washes for 30 minutes and a final 5 minute wash in 5x SSC. Embryos were mounted in ProLong Glass Antifade Mountant (Thermo Scientific).

### Image collection

Confocal images were generated using a Zeiss LSM880 with the Plan-Apochromat 25x/0.8 multi-immersion lens with oil, Plan-Apochromat 40x/1.3 oil immersion lens, or the Plan- Apochromat 63x/1.4 oil-immersion lens. Images were captured with either the internal GaAsP detector or an Airyscan detector; Airyscan processing was performed on Zen software. All images in the paper are oriented with the anterior facing the left. Image analysis was performed using Fiji^65^ and associated plugins.

### Targeted DamID

SrpC and SrpNC isoforms were cloned from pBS-KS Srp, which contains the open reading frame(ORF) for SrpC (gifted by C.Antoniewski), and the full-length SrpNC ORF was cloned from pBS-KS SrpNC, (gifted by M.Haenlin^66^) and inserted into UAS-LT3-Dam (gifted by T.Southall^33^). UAS-SrpC-Dam, UAS-SrpNC-Dam and UAS-LT3-Dam (control line) flies were crossed to hkb-Gal4 and reared at 25°C. ∼1 g of embryos were collected per condition. Embryos of the correct genotype were collected after 1 hr periods for successive lays over a 12-hr period. Embryos were aged to stage 11 and dechlorinated and fixed using standard procedures. Genomic DNA was extracted using the Qiagen QIAamp DNA Micro Kit, followed by overnight digestion at 37°C by DpnI, resulting in restriction endonuclease activity within methylated GATC sites. Digested DNA was purified using the QIAquick PCR Purification kit. Adaptors were blunt-end ligated for 2 h at 16°C using T4 DNA ligase and heat inactivated at 65°C for 20 min. The ligated DNA was then digested with DpnII to cleave any unmethylated GATC sites, and purified with a 1:1 ratio of Seramag beads. Adaptor-ligated fragments were then amplified with MyTaq and PCR purified. The amplified DNA was sonicated and digested with AlwI to remove large adaptor sequences. Custom sequencing adaptor sequences containing unique identification sequences were ligated in-place of DamID adaptor sequences, pooled and prepped for Illumina sequencing, according to previously published methods ^33^. The quality of newly generated sequencing libraries containing unique identifier sequences was assessed using a Bioanalyzer. All sequencing was performed as a single end 50 bp reads generated by NovoGene using the NovaSeq 6000. Each sub-indexed library contained 4 biological replicates, each with 2 experimental replicates of embryos collected at distinct time periods.

### Targeted DamID analysis

Sequencing data generated from UAS-SrpC-Dam and UAS-SrpNC-Dam were used to generate Log2 ratio files relative to UAS-Dam (UAS-SrpNC-Dam/UAS-Dam and UAS-SrpC- Dam/UAS-Dam) via the DamMer pipeline^37^. A False Discovery Rate of -log10(50) was used for peak calling.

### Bioinformatic analysis

Called peaks from the DamMer pipeline were mapped to nearby genes using Uropa^64^. The Motif enrichment analysis of Srp-bound peaks was performed using the FindMotifsGenome command from Homer^65^. Motif enrichment analysis in gene lists was undertaken using i- cisTarget^66,67^. Binding plots at genomic loci were created using pygenometracks^68,69^. GO term analysis was performed using the PantherGO web tool. Aggregation plots of Serpent binding profiles were made using Seqplots^70^.

### Exposure to exogenous 20E

Embryos were dechorionated, immobilized between two nylon meshes, and permeabilized by immersing the immobilized embryos in *n*-octane (Sigma) for 5 min. After evaporation of the octane, 1 ml of Schneider’s medium containing 5 × 10^−6^ M 20-hydroxyecdysone (Sigma) was applied to the double mesh for 2 h. The 20-hydroxyecdysone solution was then replaced by Schneider’s medium, and the embryos were left to develop for a further 1 hr. Control embryos received the same treatment, but without 20-hydroxyecdysone.

Embryos were laid over a 1 hr period, left to develop for 2 hrs, followed by exposure to either Schneider’s medium containing 20E or Schneider’s medium alone for a period of 2 hrs. All embryos, regardless of previous treatment, were then exposed to Schneider’s medium only for 1 hr.

### Cell segmentation and scMorphometrics

Cells were manually segmented from Z-stacks of Airyscan-processed images (0.44μm Z distance) using the interactive object drawing tool in Arivis Vision4D. To avoid variability, only the distal portion of the midgut was segmented. Each cell surface was extracted, and cells were reoriented in the apical-basal axis using Blender v4.3. Morphometric features were extracted for each cell using PyVista^71^. These features were volume, surface area, surface area:volume, cuboidness (cell volume/bounding box volume), sphericity, flatness, surface area of the most apical 10% of the cell, surface area of the most basal 90% of the cell, apical area:basal area, cell height, and cell width at the widest point. Principal component analysis of these properties was performed with Factoextra^72^.

### Fluorescence intensity quantification and cell counting

To quantify Stardust and Crumbs intensity across midguts, a 500 pixel line was drawn from the ventral end of the midgut to the dorsal end in pre-migration midguts, spanning PMECs and endoblast-derived cells. The measure tool from Fiji (RRID:SCR_025376) was used to collect intensity values which were normalised for length and to the maximum intensity value collected for each embryo.

To quantify fluorescence intensity of *lacz* FISH in different cell populations, cell outlines were traced using the draw tool in Fiji and the area outside the selection deleted. Once done in all Z slices, the segmented image was sum intensity projected and the *lacz* channel intensity recorded.

Embryos were imaged at stage 15 when intending to count EEs or AMPs. All cells were counted using the Cell Counter plugin in Fiji^62^. Pros^+^ cells were considered EEs and Snail^+^ cells were considered AMPs. Any Pros^+^Snail^+^ cells were considered a subset of EEs.

## Acknowledgements

We are thankful to the rest of the Campbell, Casali, and Strutt labs for helpful discussions. We thank Tony Southall and Robert Krautz for their advice and sharing of protocols. We thank T.Southall, , the Bloomington Stock Centre, and the Developmental Studies Hybridoma Bank for kindly sending us reagents. This work was supported by a Wellcome Trust/Royal Society Sir Henry Dale Fellowship (grant number 204615/Z/16/Z).

## Author contributions

This project was conceived by A.T.P., J.A. and K.C. Experiments were performed by A.T.P. and J.A. with contributions from Z.R. A.T.P and K.C. wrote the manuscript, which was edited and reviewed by all authors.

## Declarations of interests

The authors declare no competing interests.

**Supplementary Figure 1. Both Serpent isoforms primarily bind to intragenic regions.** (A, B) Aggregation plots show peak binding intensity between the transcription start site (TSS) and transcription end site (TES) for SrpC (green) and SrpNC (magenta). Uropa analysis shows that the majority of peaks are within gene loci for both SrpC (B) and SrpNC (D).

**Supplementary Figure 2. Serpent isoforms bind to known target genes.** TaDa binding profiles for SrpNC (magenta) and SrpC (green) at the locus for *lanb2* (A) and *gatae* (B). The y-axis is a log_2_ score for Srp binding enrichment over background binding.

**Supplementary Figure 3. Breakdown of principal components producing morphospace variability.** Plots showing which physical properties extracted from segmented cells contribute to PC1 (A) and PC2 (B). The red line represents the average expected contribution for properties. (C) A Principal Component Analysis biplot showing the distribution of cells segmented from hkb-stGFP (blue) or hkb-EcRDN (red) midguts at a post-EMT timepoint and the contribution of variables in 2D space, showing how each measured property influences each PC.

**Supplemental Figure 4.** B**l**ocking **ecdysone signalling does not affect AMP and EE specification.** Cell counts from stage 15 hkb>stGFP and hkb>EcRDN embryos for AMPs (A) and EEs (B). Black diamonds are the mean for each genotype with error bars showing standard deviation. P-values were calculated via t-test.

## Notes

### Competing Interest Statement

The authors have declared no competing interest.

### Summary of Updates

Figures 2 and 5 were revised and updated

