## Supplementary figures and images for "A hormonally regulated gating mechanism controls EMT timing to ensure progenitor cell specification occurs prior to epithelial breakdown"

### Supplemental Figure 1

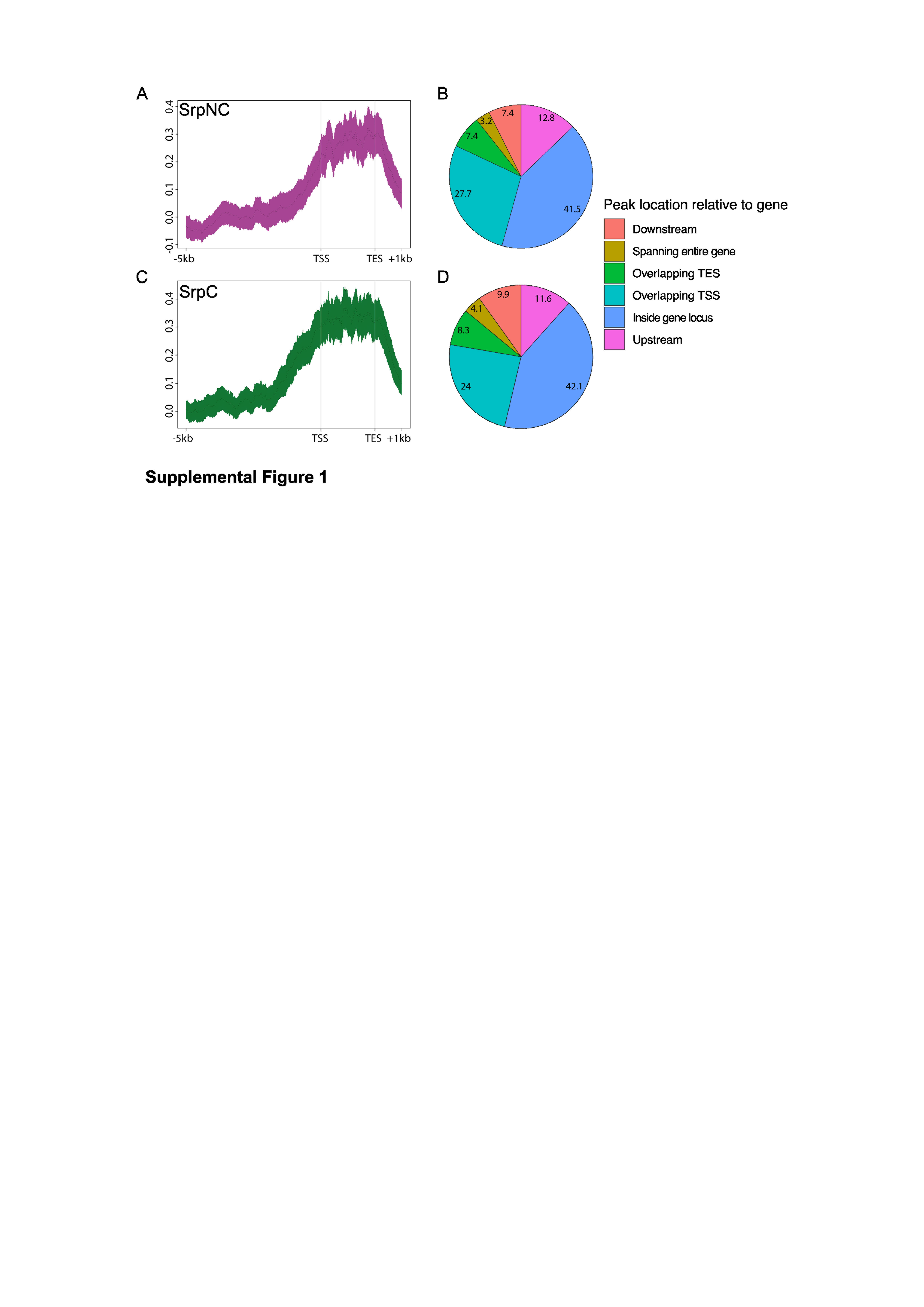

### Supplemental Figure 2

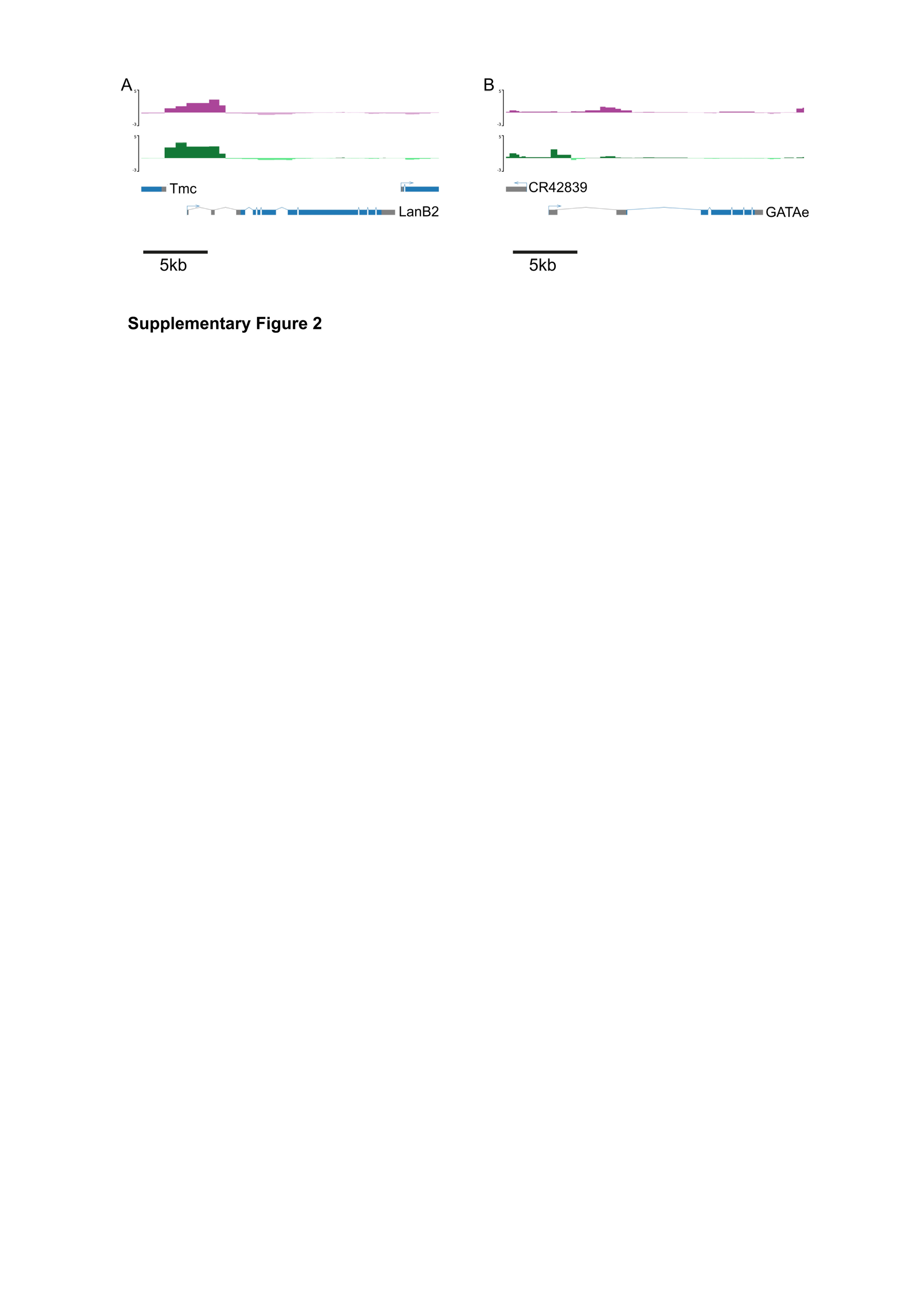

### Supplemental Figure 3

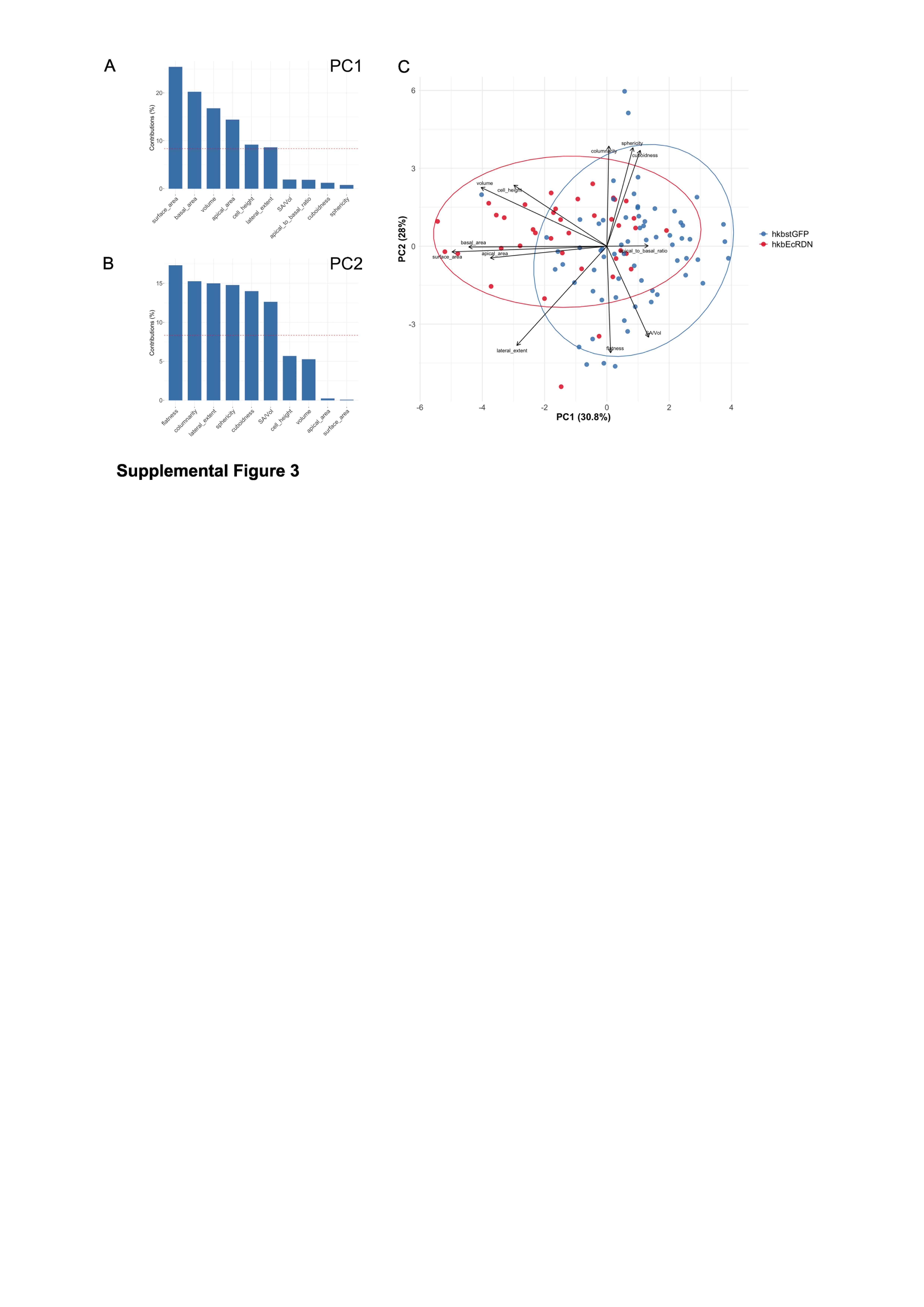

### Supplemental Figure 4

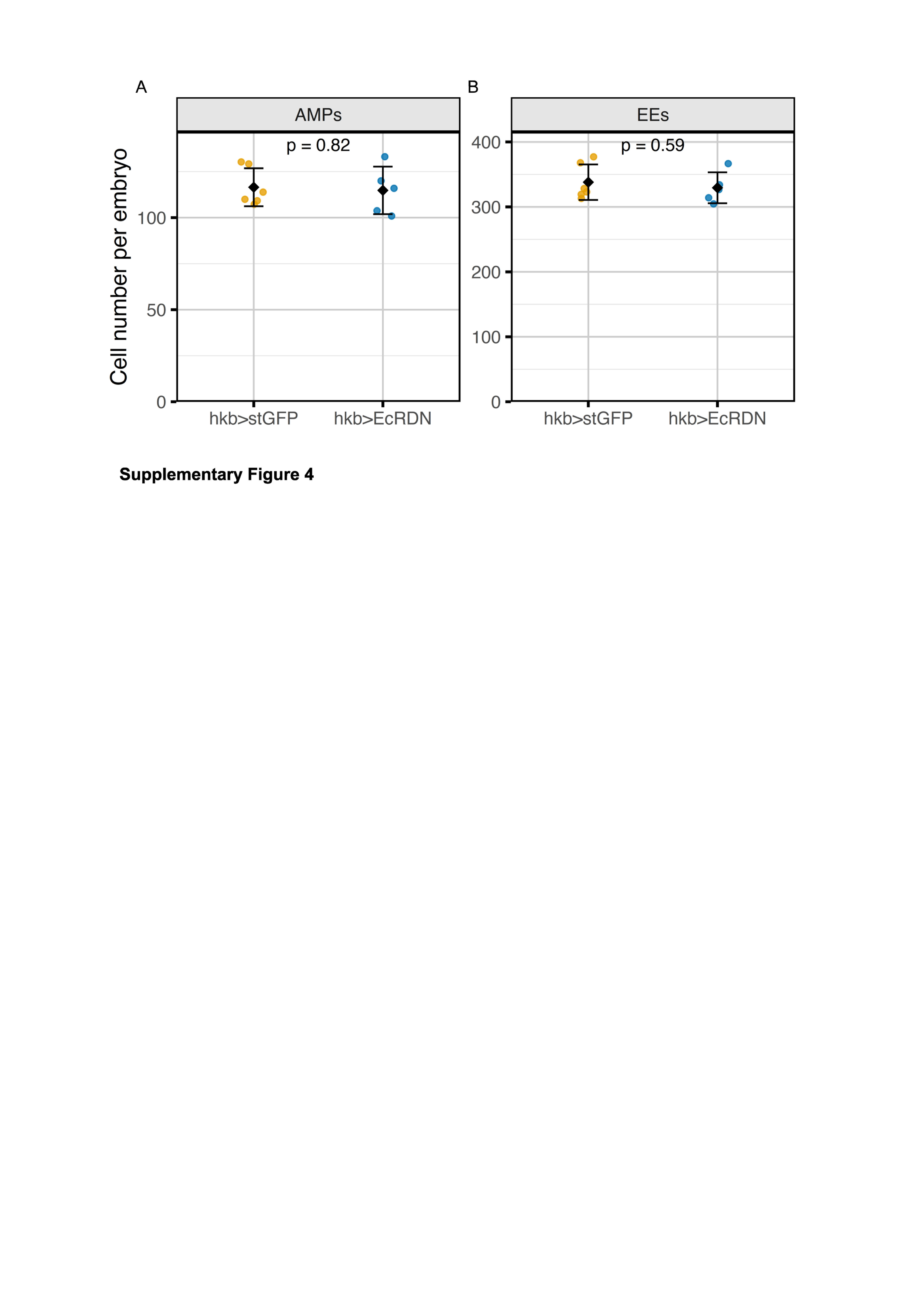
